# Structure and dynamics of human cardiac fibroblast nanotubes

**DOI:** 10.1101/2023.11.28.568871

**Authors:** S.C. Schmid-Herbstritt, G. Stief, J. Greiner, A. Felekary, J. Madl, V. Zeidler, J. Heer, P. Iaconianni, M. Koch, K. Kollmar, C. Walz, S. Nübling, T. Kok, J.R. Pronto, I. Kutschka, N. Voigt, G. Morgan, J. Dienert, T. Brox, P. Briquez, R. Peyronnet, A. Rohrbach, P. Kohl, E.A. Rog-Zielinska

**Author notes:** Correspondence to: E. Rog-Zielinska, Institute for Experimental Cardiovascular Medicine, Elsaesserstrasse 2Q, 79110 Freiburg, Germany.

## Abstract

Efficient and dynamic interactions between cardiac fibroblasts and their environment are essential for the maintenance of tissue homeostasis in healthy hearts and play an important role during pathological remodelling. Here, we investigate a relatively obscure mechanism through which human atrial fibroblasts communicate with each other, with other cells, and with the extracellular matrix (ECM) – nanotubes (NT). We investigated NT structure and dynamics in primary right atrial fibroblasts isolated from patients in sinus rhythm (SR) and atrial fibrillation (AF), in an immortalised human atrial fibroblasts cell line, and in intact human tissue, using a wide range of imaging approaches (including confocal microscopy, label-free reflection microscopy, rotating coherent scattering microscopy, and cryo-electron tomography). We show that fibroblasts maintain continuous NT activity *in vitro*, with numerous protrusions constantly probing the surrounding environment. NT structure and activity change during AF and following pharmacological (transforming growth factor-β, latrunculin B) and environmental (hypoxia) interventions. We also show that cardiac fibroblast NT mediate intercellular organelle exchange and dynamically interact with ECM. Finally, we present evidence for the presence of fibroblast-borne NT in human atrial tissue. Our results advance our understanding of how cardiac fibroblasts interact with their environment. NT are versatile structures capable of both sensory and actuating functions, and offer a dynamic and rapid communication conduit that facilitates cell–cell and cell–extracellular matrix interactions.

## Introduction

Cardiac fibroblasts support the structural, mechanical, and electrical integration of cardiomyocytes in the heart.^1^ Appropriate fibroblast activity (e.g. production and deposition of extracellular matrix [ECM] components, secretion of paracrine mediators) is necessary for the maintenance of tissue integrity and homeostasis. However, dysregulation of fibroblast behaviour can negatively affect the heart’s performance, e.g. through excessive ECM deposition. Knowledge of the mechanisms controlling the dynamic properties of fibroblasts could therefore offer a way to modulate cardiac architecture, particularly during pathological remodelling, shifting the cardiac environment to one in which appropriate electro-mechanical connectivity between various cardiac cells and ECM can be sustained.

Fibroblasts can dynamically remodel the cardiac tissue (by balancing the production, retention, or degradation of various ECM components or signalling molecules for example) in response to signals stemming from cell–cell and cell–ECM communication. Interactions between fibroblasts and other cells or the ECM are thought to occur both directly and indirectly; indirect interactions include paracrine mediators or interactions via ECM-bound surface receptors, whereas direct interactions include electrotonic coupling and formation of mechanical junctions.^1–3^ In contrast to many other cardiac cell types, fibroblasts are highly motile, able to migrate through the tissue, can dynamically ‘probe’ their environment, and rapidly form new contacts with surrounding cells and extracellular motifs. Cellular nanotubes (NT) are unique potential conduits for direct, fast, and transient cell–cell and cell–matrix communication, in part because of their structure (typically 50–600 nm wide and 5–300 µm long), allowing for dynamic communication in a confined tissue environment.^4–6^

NT exhibit high variability in terms of morphology, content, association with the substrate (attached vs ‘floating’), and potential for cell–cell material exchange. As no unambiguously specific molecular markers have been described for the various types of NT (filopodia, tunnelling nanotubes, microvilli, and cytonemes), distinguishing between them is challenging and often depends on the cell of origin and stage of NT formation. It has been proposed that filopodia may be identified by their adherence to the substrate, whereas tunnelling nanotubes in contrast ‘float’ and can establish direct cytoplasmic connections between cells.^7–11^

Regardless of classification, various NT types have been shown to be involved in many processes *in vitro*, including cell growth and migration (e.g. during embryonic development or cancer progression), apoptosis, and adhesion to ECM.^7,10^ In addition, open-ended protrusions enable homo- and hetero-cellular sharing or exchange of various small molecules (calcium, proteins, prions, antigen receptors), cellular components (genetic material, organelles), or pathogens.^12–15^ The presence of NT has been demonstrated in cultured cells of various origins (including epithelial, endothelial, mesenchymal, neuronal, immune, stem cells). Formation of NT has been shown to be increased by inflammatory signals, oxygen depletion, and metabolic stress.^16–19^ Although detecting NT in tissue is much more challenging due to their small size (often below the diffraction limit of light) and transient nature, several studies have demonstrated their presence in intact tissue.^17,20,21^

The presence of NT in cardiac cells has been described in only a handful of reports. Cardiac fibroblasts and embryonic, neonatal, and stem cell-derived cardiomyocytes have been shown to be capable of forming NT *in vitro*.^3,19,22–26^ NT were suggested to mediate alterations of the secretome, inflammasome activation, and organelle transfer. The presence of NT has also been reported in intact cardiac tissue;^3,22^ however, the dynamic behaviour and relevance of cardiac fibroblast NT and the mechanisms governing their formation and guidance in the healthy or diseased heart remain for the most part unclear.^3,19,22,25^

The number and activity of fibroblasts drastically change in the diseased heart. NT could offer a conduit for dynamic homo-or hetero-cellular communication, and may mediate the necessary adaptations of cardiac cells to changes in the tissue environment such as changes in stiffness, electrical properties, ECM content, or oxygen availability. Here, we provide a thorough characterisation of human atrial cardiac fibroblast NT *in vitro*, including their nanoscopic and microscopic morphology and motility, and their association with ECM. We also describe changes in NT properties in the context of atrial fibrillation (AF), a disease associated with structural and electrical remodelling of fibroblasts. We also explore whether pharmacological or environmental interventions can affect the properties of human cardiac fibroblast NT. Finally, we describe the presence of fibroblast NT in human atrial tissue.

## Materials and Methods

All patients gave written and informed consent prior to inclusion in the study, and investigations conformed to the principles outlined in the Declaration of Helsinki. Tissue samples were excised from the right atrial appendage as a routine procedure in the course of cannulation for extracorporeal circulation during open heart surgeries. Excised tissue samples were immediately placed in cardioplegic solution (containing: NaCl 120 mM, KCl 25 mM, HEPES 10 mM, glucose 10 mM, MgCl_2_ 1 mM; pH 7.4, 300 mOsm, room temperature) and transported to the laboratory, where they were processed by the Cardiovascular Biobank at the University Heart Center Freiburg/Bad Krozingen (approved by the ethics committee of the University of Freiburg, Nr. 607/19). Patients were either in sinus rhythm (SR), or had sustained AF (including patients with persistent, long-standing persistent, and permanent AF, defined according to ESC Guidelines).^27^ Patient demographics are listed in Table 1.

**Table 1.** Patient characteristics. All samples listed here were used for primary fibroblast isolation. Data analysed by Student’s *t*-test. Reconstr., reconstruction; repl., replacement; SD, standard deviation.

|  | **Sinus Rhythm**<br>**(SR)**<br>**N = 8** | **Atrial Fibrillation**<br>**(AF)<sup>27</sup>**<br>**N = 3** | **p-value** |
| --- | --- | --- | --- |
| Male patients, n (%) | 7 (87.5) | 3 (100) |  |
| Mean age at time of surgery, years (SD) | 64.1 (21.2) | 69.3 (6.2) | 0.71 |
| Mean BMI, kg/m<sup>2</sup> (SD) | 26.1 (5.6) | 25.6 (2.7) | 0.88 |
| Diabetes mellitus, n (%) | 1 (12.5) | 0 |  |
| Hyperlipidemia, n (%) | 4 (50.0) | 3 (100) |  |
| Arterial hypertension, n (%) | 4 (50.0) | 3 (100) |  |
| Mean systolic blood pressure, mmHg (SD) | 139.7 (26.9) | 156.7 (9.4) | 0.37 |
| Mean diastolic blood pressure, mmHg (SD) | 77.6 (20.4) | 88.3 (2.4) | 0.44 |
| Mean heart rate, bpm (SD) | 83.4 (19.4) | 62.0 (23.2) | 0.20 |
| Mean ejection fraction, % (SD) | 57.3 (2.9) | 56.3 (9.0) | 0.82 |
| Surgical procedures, n (%) |  |  |  |
| Aorto-coronary venous bypass | 2 (25.0) | 0 |  |
| Aortic valve replacement/reconstr. | 4 (50.0) | 0 |  |
| Mitral valve replacement/reconstr. | 1 (12.5) | 2 (66.7) |  |
| Tricuspid valve repl./reconstr. | 1 (12.5) | 1 (33.3) |  |
| Aortic arch replacement | 2 (25.0) | 0 |  |
| ASD closure | 1 (12.5) | 0 |  |
| Medication, n (%) |  |  |  |
| ACE inhibitors | 1 (12.5) | 1 (33.3) |  |
| ATI-receptor blocker | 2 (25.0) | 1 (33.3) |  |
| β-blocker | 2 (25.0) | 1 (33.3) | |
| Diuretics | 2 (25.0) | 2 (66.7) |  |
| Statins | 2 (25.0) | 2 (66.7) |  |
| Anticoagulants | 3 (37.5) | 3 (100) |  |
| Ca<sup>2+</sup> antagonists | 1 (12.5) | 0 |  |

In addition to samples collected in-house, human right atrial tissue from patients with AF was obtained from the University of Göttingen. Experimental protocols were approved by the ethics committee of the University Medical Center Göttingen (No. 4/11/18). Patient demographics are listed in Table 2. The tissue was chemically fixed immediately following excision in iso-osmotic Karnovsky’s fixative (2.4% sodium cacodylate, 0.75% formaldehyde, 0.75% glutaraldehyde).

**Table 2.** Patient characteristics. All samples listed here were used for intact tissue imaging. Reconstr., reconstruction; SD, standard deviation.

|  | **Atrial Fibrillation (AF)<sup>27</sup>**<br>**N = 3** |
| --- | --- |
| Male patients, n (%) | 1 (33.3) |
| Mean age at time of surgery, years (SD) | 72.0 (4.3) |
| Mean BMI, kg/m<sup>2</sup> (SD) | 26.9 (5.1) |
| Diabetes mellitus, n (%) | 1 (33.3) |
| Hyperlipidemia, n (%) | 1 (33.3) |
| Myocardial infarction, n (%) | 1 (33.3) |
| Arterial hypertension, n (%) | 2 (66.7) |
| Mean heart rate, bpm (SD) | 82 (7.1) |
| Mean ejection fraction, % (SD) | 60.7 (4.9) |
| Surgical procedures, n (%) |  |
| Aorto-coronary venous bypass | 2 (66.7) |
| Aortic valve replacement/reconstr. | 1 (33.3) |
| Mitral valve replacement/reconstr. | 1 (33.3) |
| Medication, n (%) |  |
| ACE inhibitors | 2 (66.7) |
| ATI-receptor blocker | 1 (33.3) |
| β-blocker | 3 (100) |
| Diuretics | 1 (33.3) |
| Statins | 3 (100) |
| Anticoagulants | 2 (66.7) |
| Ca<sup>2+</sup> antagonists | 1 (33.3) |

### Fibroblast isolation, culture, and pharmacological treatment

Primary patient-derived atrial fibroblasts were obtained using the outgrowth method as described previously.^28^ Only cells from passages 1–4 were used for experiments. In addition to primary atrial fibroblasts, an immortalised human atrial fibroblast (HAF) cell line was used for a subset of experiments.^29^ All cells were cultured in Dulbecco’s modified Eagle medium (Thermo Fisher, Waltham, MA, USA), supplemented with 10% fetal calf serum, 1% penicillin-streptomycin, 1× non-essential amino acids, and 0.5 mM L-ascorbic acid 2-phosphate sesquimagnesium salt hydrate (all Sigma-Aldrich, St. Louis, MO, USA). Cells were maintained at 37°C in an atmosphere of air supplemented with 5% CO_2_. Cells were passaged when 70-80% confluence was reached. Prior to experiments, cells were plated on 100 µg/mL laminin-coated dishes (see relevant sections for details). Seeding densities are indicated where appropriate.

In addition to standard 2D cultures, cells were seeded in 3D matrices composed of Matrigel (Beckton Dickinson, Franklin Lakes, NJ, USA) at a density of 15,000 cells per 150 µl Matrigel, and cultured for 7 days prior to imaging. Pharmacological treatment was carried out with 10 ng/mL transforming growth factor-β (TGF-β1; Abcam, Cambridge, UK) for 24 hours, or 60 nM latrunculin B (Tocris, Bristol, UK) for 2 h prior to experiments. Control samples were treated with vehicle (ethanol, dilution 1:250,000) alone.

### Fluorescence staining

For imaging of fixed cells, cells were seeded on glass coverslips placed in 12-well plates at a density of 5,000 cells/well. Cells were chemically fixed 3 days after seeding using 2% formaldehyde, at room temperature, for 15 minutes. Cells were then washed with phosphate-buffered saline (PBS). Permeabilisation and blocking of non-specific antibody binding were carried out using PBS supplemented with 5% fetal calf serum, 2.5% bovine serum albumin, and 0.1% Triton X-100, for 30 minutes at room temperature and with agitation. Immuno-staining was conducted using mouse anti-β-tubulin 1E1-E8-H4 antibody (dilution 1:300, cat. nr. ab131205, Abcam, Cambridge, UK), or goat anti-collagen Iα1 antibody (dilution 1:20, cat. nr. MBS316282, MyBioSource, San Diego, CA, USA) in PBS supplemented with 2.5% fetal calf serum and 1% bovine serum albumin, overnight at 37°C and with agitation. After washing, the cells were incubated with species-appropriate fluorophore-conjugated secondary antibodies (all Thermo Fisher, Waltham, MA, USA). Fixed cells were additionally stained with membrane marker CellBrite (dilution 1:200, Biotium, Fremont, CA, USA) and actin marker Phalloidin (dilution 1:1000, Abcam, Cambridge, UK) according to the manufacturers’ instructions. Following staining, the samples were mounted on glass slides using PermaFluor mounting medium (Thermo Fisher, Waltham, MA, USA), sealed with nail polish, and kept in the dark until imaging. Images (length, actin content, number of NT) were analysed manually using a custom Python script and Napari visualisation tool (https://napari.org/).^30^ Script is available upon request.

For live-cell imaging, cells were seeded on 35 mm ibidi glass-bottom dishes (ibidi, Gräfelfing, Germany) at a density of 15,000 cells/well, and imaged on day 3 after seeding. The cell membranes were visualised using Cell Mask dye (dilution 1:1000, Thermo Fisher, Waltham, MA, USA), and cell volume with cytoplasm marker CellTracker (dilution 1:200, Thermo Fisher, Waltham, MA, USA), according to the manufacturers’ instructions. Mitochondria were visualised using MitoView dye (100 nM, Biotium, Fremont, CA, USA). Collagen in live cultures was visualised using goat anti-collagen Iα1 antibody (dilution 1:20, cat. nr. MBS316282, MyBioSource, San Diego, CA, USA), conjugated to CF555 (Mix-n-Stain CF Antibody Labeling Kit, Biotium, Fremont, CA, USA)

### Confocal fluorescence and reflection microscopy imaging

Image acquisition was carried out using a confocal laser scanning microscope (Leica TCS SP8 X, Leica Microsystems, Wetzlar, Germany) and HC PL APO 40x/1.1 WATER or HC PL APO CS2 63×/1.4 OIL objectives. The system was controlled using the Leica Application Suite X (LAS-X). Samples were illuminated using appropriate wavelengths, which were delivered from a white light laser.

Live-cell imaging was carried out using a combination of fluorescence and reflection microscopy. Reflection imaging was conducted using 5 laser lines simultaneously (600 nm, 615 nm, 630 nm, 645 nm, 660 nm), with acousto-optic beam splitter operated in reflection configuration. Signal was detected within a 590 nm–671 nm wavelength window. The cells were imaged in a stage-top incubator (Tokai HiT, Gendoji-cho, Fujinomiya-shi. Shizuoka-ken, Japan), at 37°C, in a humidified atmosphere of air supplemented with 5% CO_2_. Hypoxia was implemented by lowering the oxygen levels to 1% (substitution with nitrogen). For assessment of NT motility, sequences of images were acquired over a period of 15 minutes at 4 frames/minute, with 5 to 7 sequences per sample (primary fibroblasts), or 12 frames/minute, with 3 to 4 sequences per sample (HAF cells). For assessment of mitochondria exchange, the time series were acquired at 30 frames/hour. Representative images were denoised using SUPPORT denoising software.^31^

### Deep Learning-based NT tip tracking

NT motility was assessed using an automated pipeline that utilises deep learning for NT tip detection and a linear assignment problem (LAP) tracker to link detected tips. We trained a fully convolutional neural network using nnU-Net.^32^ The training dataset consisted of 35 time series (2048 x 2048 pixels, 62 frames total), of which 33 were sparsely annotated, and two were densely annotated. In the sparsely annotated time series, a fraction of the tips and their immediate surrounding (10-pixel radius) were annotated using TrackMate^33^ plugin within the Fiji software^34^ (https://imagej.net/software/fiji/; see Supplemental Fig. S1). The 3D U-Net variant of nnU-Net was trained using default parameters (1000 epochs, 5-fold cross-validation) and evaluated on one densely annotated test set. The average dice coefficients were 0.88, 0.66, and 0.64, for training, validation, and test, respectively (see Supplemental Fig. S2). The detections were processed by the Python wrapper ‘PyImageJ’^35^ of ImageJ and TrackMate: the tip probabilities were processed by the TrackMate’s MaskDetector and then tracked using the ‘Sparse LAP Tracker’ (allowed frame gap: 1, maximal gap closing distance: 3 frames, maximum linking distance: 3 frames). Tracks with a duration of 45 seconds or less were excluded from the analysis. Using TrackMate, the mean tip speed, and mean directional change rate were calculated. The tracking was evaluated on the test set and reached an F1 score of 0.76 (see Supplemental Fig. S3). In the test set, mean error of mean tip speed was 0.0021 µm/s (7.95% error), and mean error of mean directional change rate was 0.0061 radian (7.36% error). All automatically analysed stacks (SR N = 57, AF N = 86, HAF N = 15) were visually inspected for errors, and stacks with corrupted image quality were excluded (see Supplemental Fig. S4).

### Rotating coherent scattering microscopy

Cells were seeded on 35 mm ibidi glass-bottom dishes (ibidi, Gräfelfing, Germany) at a density of 25,000 cells/well. Imaging was carried out on day 3 after seeding at the Department of Microsystems Engineering of the University of Freiburg, using label-free super-resolution rotating coherent scattering microscopy.^36,37^ Imaging was performed in dark-field mode, allowing us to yield higher contrast compared to bright-field mode. Since NT typically hover above the substrate, we also employed non-total internal reflection mode allowing for imaging further away from the glass surface. The microscope employed a rotating focused laser beam in the back focal plane of an objective (α Plan-Apochromat 63×/1.46 OIL; Zeiss, Oberkochen, Germany), with three different colour laser beams combined in a laser combiner box (C-FLEX laser beam combiner, Huebner Photonics, Kassel, Germany). Scattered light was captured by a high-speed camera (Oryx 10GigE, Sony IMX252 Color; Teledyne FLIR, Wilsonville, Oregon, USA). Samples were illuminated with a 473 nm laser at 1 mW power. The cells were imaged in a stage-top incubator (ibidi, Gräfelfing, Germany), at 37°C, in a humidified atmosphere of air supplemented with 5% CO_2_. Sequences of images were acquired over a period of 4 seconds at 50 frames/second, with 25 to 30 sequences per sample. NT width was measured in the first snapshot of the time sequence as full width at half maximum in a line profile, using Fiji software.^34^ NT fluctuations were measured as (full width at half maximum in a time projection of whole series/full width at half maximum in the first snapshot of the time series), as described previously.^36^

### Scanning electron microscopy

Cells were seeded on glass coverslips placed in 12-well plates at a density of 5,000 cells/well. On day three after seeding the samples were fixed with iso-osmotic Karnovsky’s fixative (2.4% sodium cacodylate, 0.75% formaldehyde, 0.75% glutaraldehyde) for 10 minutes at room temperature, and washed with 0.1 M sodium cacodylate buffer. Dehydration was carried out with increasing concentrations of ethanol (10%, 25%, 50%, 75%, 90%, 95%, 100%) in steps of 15 minutes. Samples were then incubated with 50% hexamethyldisilazane in ethanol for 30 minutes, and finally placed in 100% hexamethyldisilazane and left to air-dry overnight. The coverslips were mounted onto SEM stubs and sputtered with platinum (8 nm thickness). Images were acquired with a Quattro SEM microscope (Thermo Fischer, Waltham, MA, USA) at the Institute for Disease Modeling and Targeted Medicine (IMITATE) of the University of Freiburg.

### Transmission electron microscopy

For cryo-electron tomography (cryo-ET), the cells were seeded on Quantifoil R 2/2 200 gold mesh grids (Quantifoil, Großlöbichau, Germany) coated with 100 mg/mL poly-L-lysine and 50 µg/mL collagen I, at a density of 1,000 cells/grid. On day 1 after seeding the grids were incubated with 10 nm colloidal gold particles for 1 minute and rapidly frozen in liquid ethane using GP2 plunge-freezer (Leica Microsystems, Vienna, Austria). Tilt series of vitrified samples were collected using a Tecnai F20 microscope (FEI Company, now Thermo-Fisher Scientific, Eindhoven, The Netherlands) at the CU Boulder Center for Cryo Electron Tomography at the University of Colorado, Boulder. IMOD software was used for volume reconstruction.^38^

For room temperature-ET, the cells were seeded on 6 mm sapphire disc (Wohlwend, Sennwald, Switzerland) at a density of 450 cells/disc. On day 3 after seeding the samples were either chemically fixed with iso-osmotic Karnovsky’s fixative (2.4% sodium cacodylate, 0.75% formaldehyde, 0.75% glutaraldehyde) or high-pressure frozen between copper planchettes using EM ICE (Leica Microsystems, Vienna, Austria). Vitrified samples were freeze-substituted in 1% osmium tetroxide/0.1% uranyl acetate in acetone (AFS2, Leica Microsystems, Vienna, Austria), and processed to Epon-Araldite resin as previously described.^39^ Tissue samples were stained with 1% osmium tetroxide, and processed to Epon-Araldite resin as previously described.^40^ Tilt series of semi-thick (300 nm) sections were imaged using a Tecnai F30 microscope (FEI Company, now Thermo-Fisher Scientific, Eindhoven, The Netherlands) at the Electron Microscopy Core Facility at the European Molecular Biology Laboratory (EMBL) in Heidelberg.^40^ IMOD software was used for image reconstruction.^38^

### Data analysis

Data are presented as violin plots (showing distribution, median, and quartiles). Data visualisation was performed using Prism 9 (GraphPad, San Diego, CA, USA). Where possible, data were fitted with linear mixed-effects models, with the grouping variable ‘Patient’ or ‘PatientNumber’, ‘Condition’, and ‘Treatment’ as predictor variables. The comparison of ‘lat B’ and ‘TGF-β’ was not tested. Two models, one with an interaction term between the predictor variables, and one without an interaction term, were compared with an F-test, and the model that showed an improvement in the fit to the data was chosen. An ANOVA was then performed on the linear mixed-effects model and the p-values were corrected with the Bonferroni-Holm method. This process used the ‘fitlme’, ‘coeftest’ and ‘ANOVA’ functions from MATLAB’s Statistics and Machine Learning Toolbox (Natick, MA, USA) and the ‘fwer-holmbonf’ function.^41^ A subset of data (Fig. 3) was analysed using the nested one-way ANOVA function of Prism 9 (GraphPad, San Diego, CA, USA). P<0.05 was taken to indicate a statistically significant difference between groups.

## Results

### Morphology and properties of human atrial fibroblast NT in vitro

All cardiac fibroblasts derived from human right atrial appendages (both SR and AF) contained numerous NT *in vitro*, as visualised by confocal and scanning electron microscopy (Fig. 1A, B). The majority of NT contained actin (Fig. 1C).

**Figure 1.**
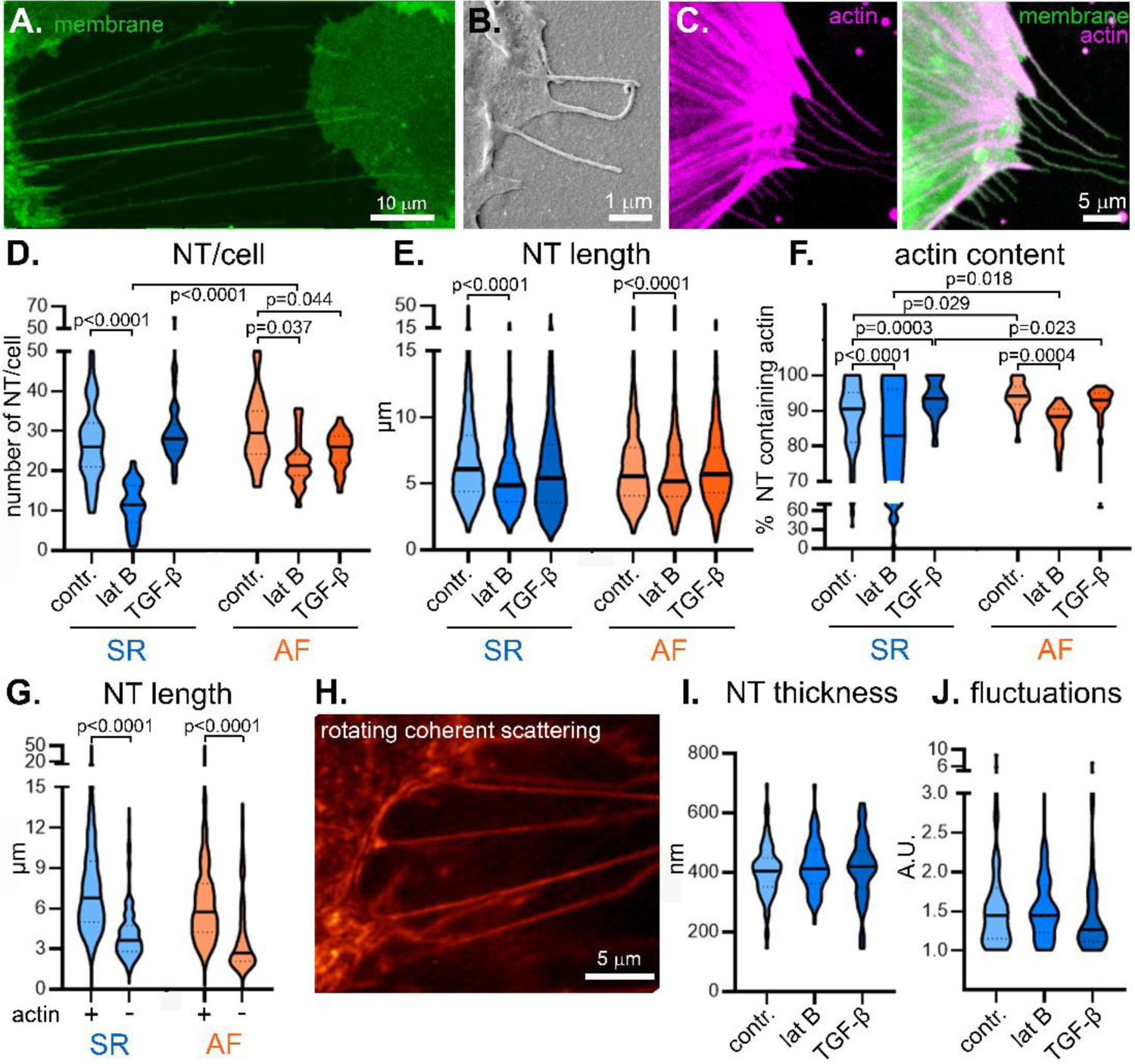
Structural properties of human primary cardiac fibroblast-borne NT and their modulation using pharmacological interventions in vitro. A–C: NT were imaged using confocal microscopy (A: live cells, fluorescent membrane dye; C: chemically fixed cells, fluorescent actin and membrane dyes), and scanning electron microscopy (B). D–F: Number (D) and length (E) of NT and percentage of NT containing actin (F) in SR and AF samples, under control conditions (contr.) and following treatment with latrunculin B (lat B) and TGF-β. Measured using confocal microscopy images of chemically fixed cells (as in C). G: Comparison of the length of NT either containing or not containing actin in control SR and AF samples. Measured using confocal microscopy images of chemically fixed cells (as in C). H: Live human primary cardiac fibroblasts from patients in SR were imaged using rotating coherent scattering microscopy. I, J: NT thickness (I) and fluctuations (J, fluctuations calculated as NT width in a time projection over 4 seconds/ NT width in the first frame), measured using data acquired by rotating coherent scattering microscopy (as in H). N = 4 SR and 3 AF patients, 282 and 202 cells, 7621 and 5246 NT (D–G); N = 3 SR patients, 86 cells, 621 NT (I, J).

The number and length of NT were not statistically different between SR and AF control fibroblasts (Fig. 1D, E). The percentage of NT containing actin was higher in AF than in SR fibroblasts. Treatment with latrunculin B decreased the number, length, and the percentage of NT containing actin in both SR and AF fibroblasts (Fig. 1D–F). Treatment with TGF-β decreased the number of NT in AF fibroblasts, but not SR fibroblasts (Fig. 1D). TGF-β increased the percentage of NT containing actin in SR fibroblasts, but not AF fibroblasts (Fig. 1F). Actin content was positively correlated with the NT length in both SR and AF control samples (Fig. 1G). The pharmacological treatments had no statistically significant effect on NT thickness or NT fluctuations, the latter being a surrogate measure of stiffness (Fig. 1H–J).

### Dynamic behaviour of NT in vitro

NT tip movements were tracked in reflection microscopy time-lapse datasets (Fig. 2A). NT tips in AF fibroblasts were slower compared to SR fibroblasts. Latrunculin B caused a decrease in NT tip velocity in SR fibroblasts, but had no statistically significant effect on AF fibroblasts (Fig. 2B). TGF-β caused an increase in NT tip velocity in AF fibroblasts. The mean directional change, reflecting the degree to which NT tips changed direction during the duration of the track, was not statistically different between SR and AF fibroblasts, and was not affected by latrunculin B or TGF-β treatments (Fig. 2C).

**Figure 2.**
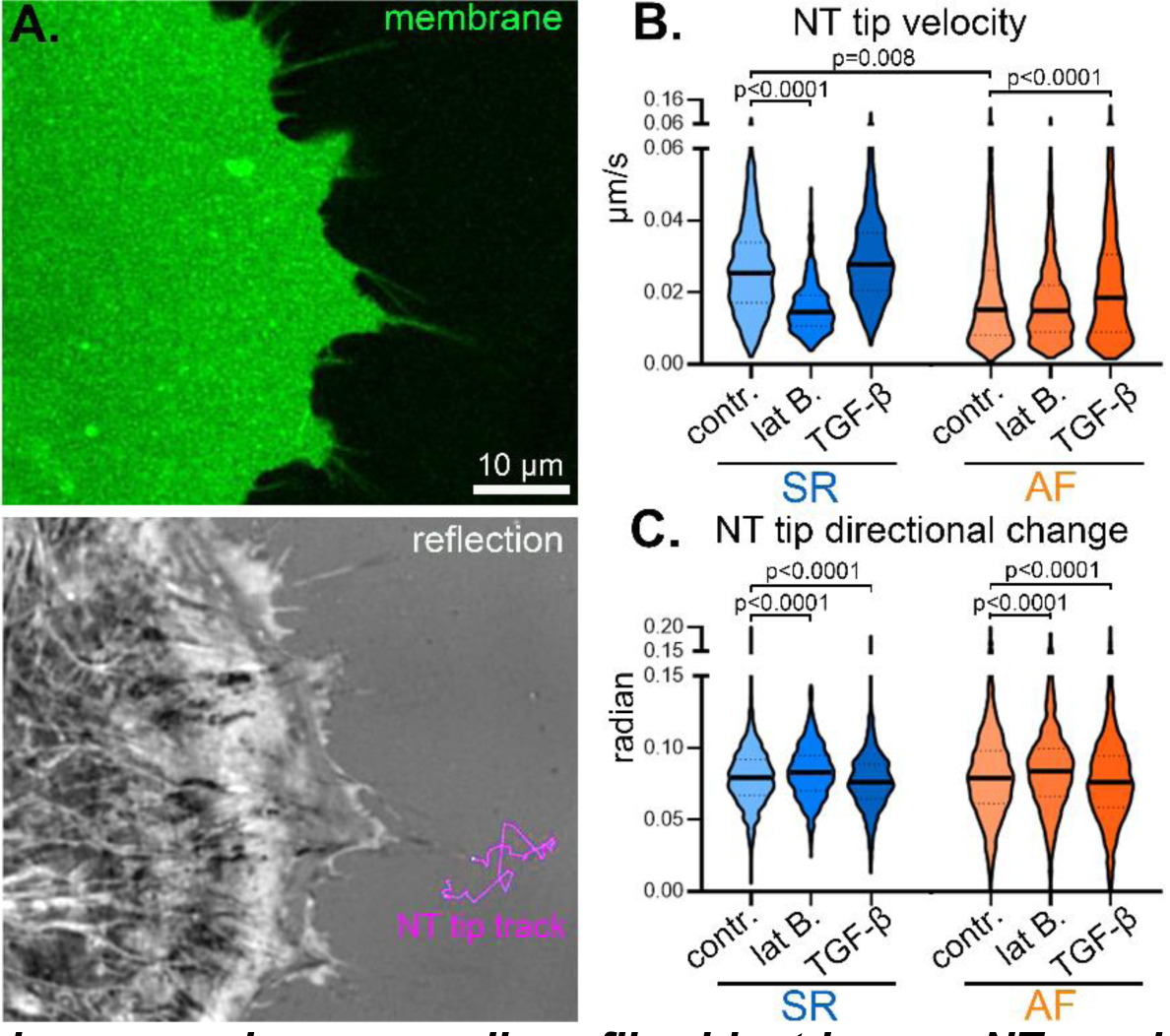
Motility of human primary cardiac fibroblast-borne NT and its modulation using pharmacological interventions in vitro. **A:** A representative image of a primary human atrial fibroblast imaged using confocal microscopy in the presence of fluorescent membrane dye (top) and using label-free time-lapse reflection microscopy (bottom). Example NT tip progression (‘track’) over 10.5 minutes indicated in magenta. Each track is composed of lines drawn between the sequential locations of the tip. **B, C:** NT tip velocity (B) and mean directional change (C), a measure of the average angle between each sequential pair of lines over an entire track; a value of 0 indicates a straight-line track) in SR and AF samples, and following pharmacological interventions. N = 4 SR and 3 AF patients, 145 and 136 cells, 12713 and 5489 NT.

To assess the effect of hypoxia on NT, we performed label-free reflection microscopy imaging under environmental control, subjecting immortalised HAF cells (Fig. 3A) to normoxia (30-90 minutes) followed by hypoxia (1% oxygen, for 30-90 minutes). The motility of NT was assessed during normoxia and after 30 minutes of hypoxia. Hypoxic conditions caused an increase in the number of NT (Fig. 3B), with no statistically significant effect on NT tip velocity or mean directional change (Fig. 3C, D).

**Figure 3.**
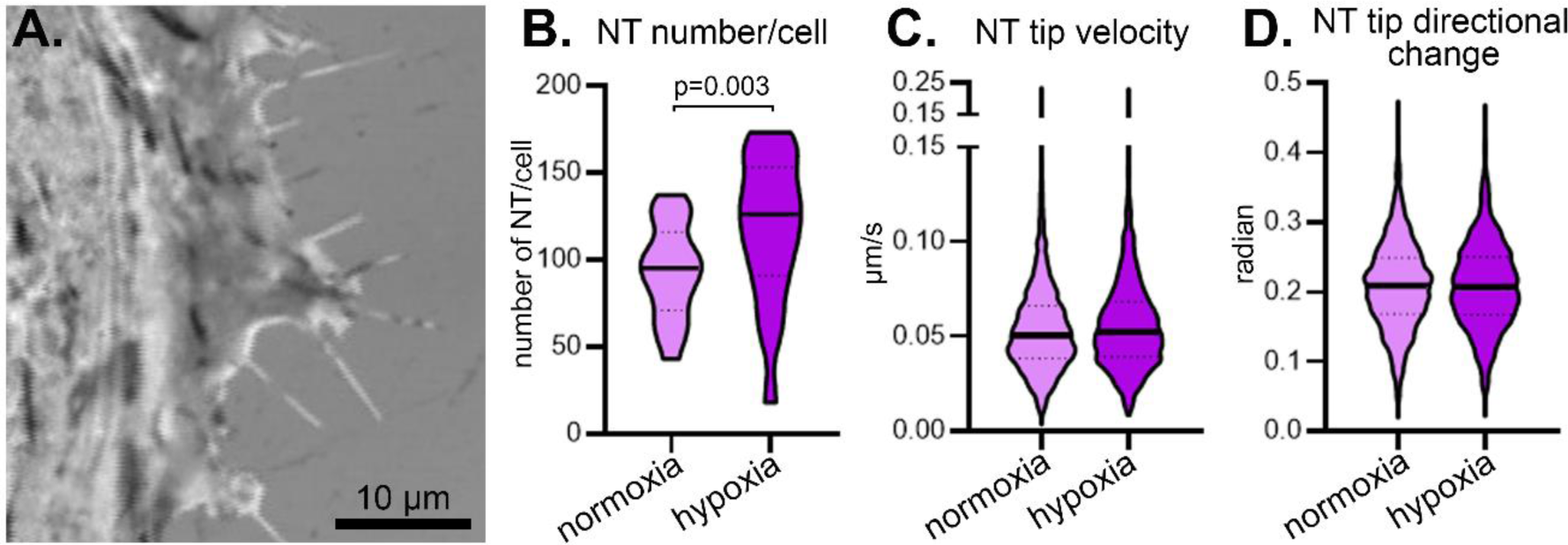
Impact of hypoxia on immortalised human atrial fibroblasts (HAF). **A:** A representative image of a HAF cell acquired using reflection microscopy. Notably, immortalised fibroblasts under control conditions contained more NT than primary human atrial fibroblasts from patients in SR used in other experiments (SR primary fibroblasts: 28 NT/cell ± 15; HAF: 94 NT/cell ± 26; average ± standard deviation; p<0.001, Student’s t-test, N = 96 primary fibroblasts and 19 HAF cells). **B:** The number of NT in HAF cells increased following exposure to hypoxia. **C, D:** NT tip velocity (C) and mean directional change (D), a measure of the average angle between each sequential pair of lines over an entire track (a value of 0 indicates a straight-line track), in HAF cells exposed to hypoxia was not statistically different when compared to normoxic conditions. N = 2 independent experiments, 19 cells (B); N = 2 independent experiments, 77-81 cells, 3784 NT (C, D).

### Nanostructure and content on NT in vitro

Cryo-ET allowed for closer examination of the structure and content of NT in immortalised HAF cells. NT contained actin fibres with the typical helical structure (Fig. 4A). NT assumed diverse geometries, including straight tubes or ‘pearls on a string’ (Fig. 4B). NT often contained vesicles or vesicle-enclosed organelles (Fig. 4B, C). Finally, we often observed bundles composed of multiple NT (Fig. 4D). NT containing microtubules were only rarely observed (Supplemental Fig. S5).

**Figure 4.**
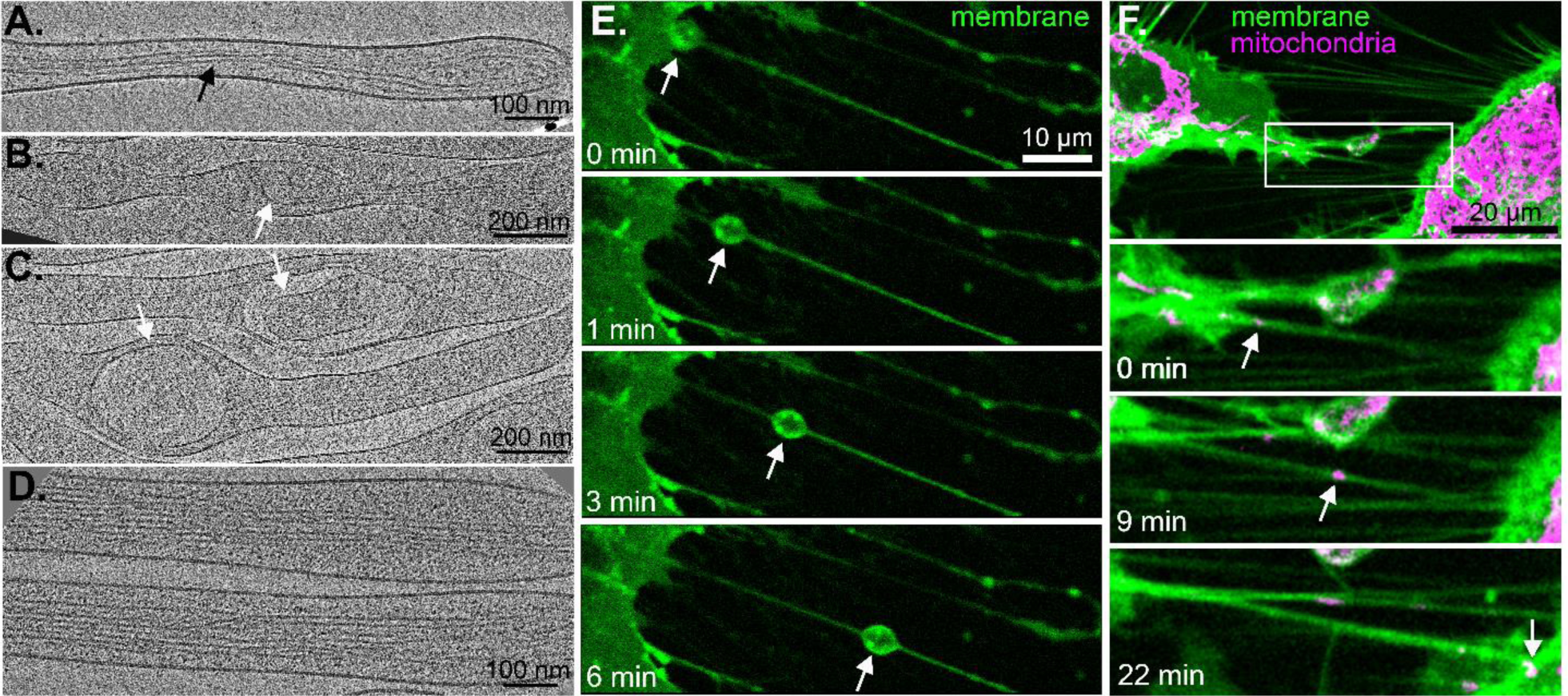
Nanostructure of NT and NT-mediated intercellular content exchange. **A–D:** NT in HAF cells contained numerous actin fibres (A–D, black arrow), vesicles (B, C; white arrow), and vesicle-enclosed organelles (C, white arrows), as visualised by cryo-ET. **E, F:** NT-mediated content exchange (white arrows) between two individual fibroblasts visualised using time-lapse confocal microscopy in the presence of fluorescent membrane (E, F) and mitochondria dyes (F).

Transport of cargo along NT was visualised using time-lapse confocal microscopy in the presence of fluorescent membrane or mitochondria dyes, using immortalised HAF cells. We observed frequent exchange of ikely vesicular cargo (Fig. 4E) and of mitochondria (Fig. 4F) along NT connecting two individual cells. The average rate of transport of cargo was 2.68 µm/minute (± 1.03, standard deviation; N = 14 events).

### Association of NT and ECM in vitro

NT were often seen in close direct contact with fibrous ECM components such as collagen I (Fig. 5A, B). Time-lapse imaging of HAF NT in the presence of fluorophore-conjugated anti-collagen I antibodies demonstrated that NT dynamically interact with collagen deposits and are able to change the arrangement and placement of fibres through direct physical interactions, such as pulling or dragging (Fig. 5C).

**Figure 5.**
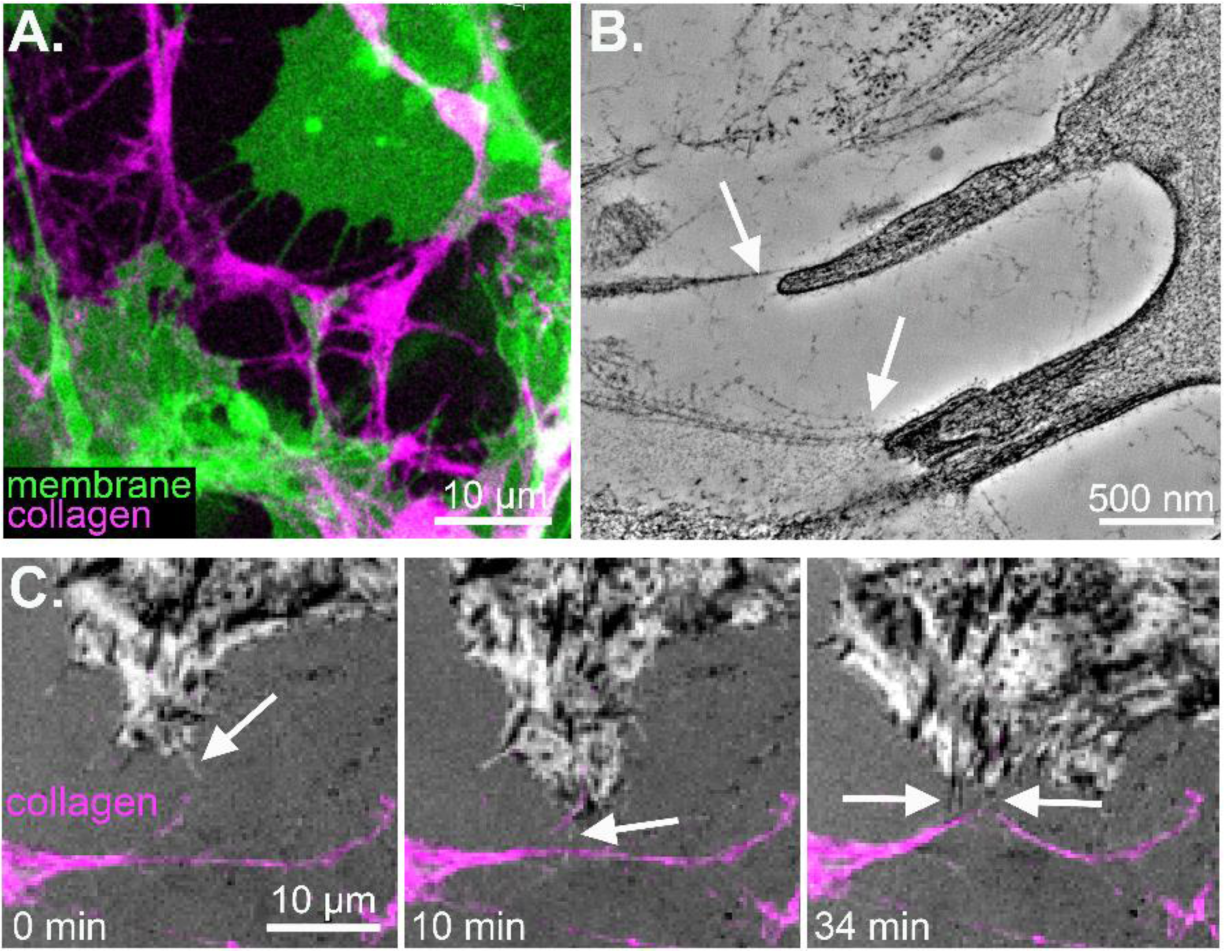
Association of NT with ECM components. **A:** HAF cell-derived NT were found to directly associate with ECM components such as collagen I, identified using confocal microscopy of chemically-fixed fibroblasts, fluorescent membrane dye, and antibodies against collagen I. **B:** Room temperature-ET of HAF cells demonstrated the association of NT and fibrous ECM components (white arrows). **C:** Time-lapse combined reflection and fluorescence microscopy in the presence of fluorophore-conjugated antibodies against collagen I demonstrated the dynamic interaction of NT with collagen I deposits (white arrows) in HAF cell cultures.

### Fibroblast NT in intact human atria and in 3D culture

Fibroblast NT-like structures were seen in intact human AF tissue, as visualised using room temperature-ET of chemically fixed samples (Fig. 6A). The apparent morphology of tissue fibroblasts was comparable to the architecture of fibroblasts when cultured in 3D matrices *in vitro* (Fig. 6B). Tissue fibroblast NT were seen in close proximity to other cell types (e.g. cardiomyocytes) and were frequently associated with ECM components (Fig. 6C).

**Figure 6.**
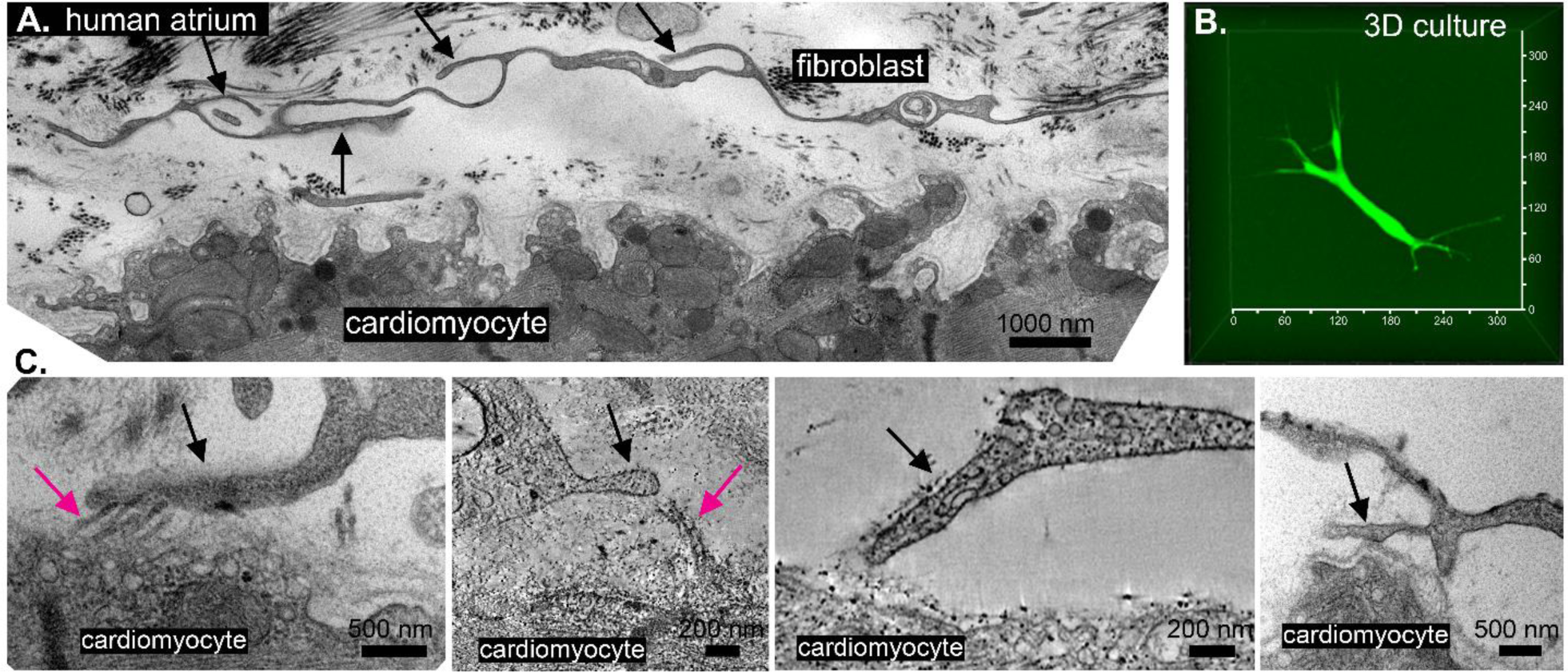
NT-like structures in intact human tissue and 3D cultures. **A:** Fibroblast-derived NT-like structures (black arrows) were seen in samples obtained from patients in AF, imaged using room temperature-ET. **B:** Morphology of a primary human SR atrial fibroblast cultured in a 3D matrix (Matrigel), and visualised using a cytoplasmic dye. **C:** Human AF tissue NT (black arrows) were often seen in close proximity to cardiomyocytes and fibrous ECM components (magenta arrows). Representative ET images from N = 3 AF patients.

## Discussion

Overall, we have demonstrated that virtually all cardiac fibroblasts derived from human atria contain numerous NT. NT activity changes during disease, and both NT structure and motility can be modulated by pharmacological and environmental factors. NT interact dynamically with collagen deposits *in vitro*, suggesting that cardiac fibroblasts perform some of their roles controlling ECM architecture via NT. Finally, we provided evidence for the presence of fibroblast-derived NT-like structures in intact human atria, and their association with other cell types and ECM deposits.

NT are extremely labile and fragile structures, and this – combined with the fact that their diameter is often below the limit of light diffraction – makes imaging of NT difficult. We used a combination of confocal fluorescent microscopy, label-free reflection microscopy (allowing us to mitigate the negative effects of the membrane dyes or strong laser exposure), super-resolution rotating coherent scattering microscopy,^36^ scanning electron microscopy, and cryo- and room temperature ET^42^ to characterise NT morphology and properties.

Human atrial fibroblast-derived NT share many of the properties reported for NT from other organ systems, including their length, width, and the fact that most contain actin.^7,10,21,24^ NT nanostructure was very heterogeneous – NT assumed different shapes and varied in content – suggesting the existence of NT sub-populations which may plausibly have different properties, lifespans, content, and roles.^43^ In order to assess whether NT morphology or behaviour differs between healthy and diseased human cardiac fibroblasts, we used fibroblasts derived from right atrial appendage obtained from patients in SR or suffering from AF, a disease characterised by severe remodelling, including changes in the electrical (ion channel function) and mechanical (ECM accumulation) properties of the tissue.^44^ In order to mimic pathological tissue states, we used pharmacological treatment with TGF-β, a known paracrine modulator shown to mediate fibroblast activation and ECM production in the injured heart. We also used latrunculin B, an actin disruptor, to correlate changes in NT content and behaviour with their structural and functional properties. Finally, as cardiac dysfunction is often associated with a reduction in the availability of oxygen, we assessed cardiac fibroblast NT during hypoxia to mimic tissue ischaemia.

As previously demonstrated in other studies,^13^ actin appeared to be a crucial component of NT, with the length of individual NT strongly correlated with their actin content. This suggests that actin fibres play an important role in NT maintenance or elongation. Indeed, the formation of NT (initiation, elongation, and stabilisation) has been shown to involve many actin regulators, such as Rho GTPases, I-BAR proteins, actin nucleators, actin bundlers, and motor proteins.^6,13^

Interestingly, NT in fibroblasts derived from AF tissue were more likely to contain actin when compared to SR cells. Actin disruption caused a decrease in the number and length of NT and the percentage of actin-containing NT in both SR and AF fibroblasts. The effects of latrunculin B on NT number or actin content were less pronounced in AF when compared to SR fibroblasts, suggesting that pathological NT remodelling in AF is associated with changes in the properties of NT actin core, possibly as a consequence of excessive fibroblast activation – this is supported by the observation that TGF-β treatment of SR fibroblasts increased the percentage of NT that contained actin to the same levels seen in AF fibroblasts, with no effect on AF NT. The effects of actin disruption or TGF-β were not reflected by any changes in the surrogate NT stiffness measure (NT fluctuations).

NT in AF fibroblasts demonstrated slower tip velocity when compared to SR fibroblasts. The exact consequences of lower NT motility in AF are currently unclear, but it is plausible that less motile NT might be unable to support sufficiently quick NT-based sensing and communication during pathological remodelling. Actin disruption caused a decrease in NT tip velocity in SR fibroblasts, indicating that actin may play a role in determining NT motility. Actin disruption had no effect on NT tip velocity in AF fibroblasts. In contrast, TGF-β caused an increase in NT tip velocity in AF but not SR samples – although tip velocity in AF upon TGF-β did not increase to the levels seen in SR cells, however. Actin disruption increased, and TGF-β decreased, the mean directional change (an indication of the NT ‘searching’ behaviour) in both SR and AF fibroblasts, suggesting that both actin and FB activation state are important determinants of the NT tip progression linearity.

While the presence of microtubules in NT was observed, it was rare and usually only seen in thicker NT. It is plausible that the incorporation of microtubules could increase the rigidity of NT and prolong the lifespan of NT,^7^ but the exact role of microtubules in possibly aiding NT reinforcement, or NT-aided intercellular interaction or transport, is currently unclear. An additional mechanism through which NT could be reinforced is bundling. Consistent with previous studies,^45^ we have observed that NT often form multi-NT bundles, potentially lending additional stability to the conduit.

Subjecting fibroblasts to hypoxia caused an increase in the number of NT, consistent with previous studies in other cell types.^19,46^ It is plausible that acutely decreased oxygen availability could create an environment in which intercellular communication would rise in importance and become a determinant of the survival of more sensitive cells (e.g. cardiomyocytes in the heart). In previous studies, hypoxia triggered fibroblasts to ‘donate’ mitochondria to ‘stressed’ cells, possibly to alleviate the metabolic crisis, restore cell function, and prevent massive recipient cell loss.^24,26,47–53^ In the current study, we observed intercellular transfer of cellular material (likely vesicles and mitochondria). Transport of cargo along NT occurred at relatively high speeds (2.68 µm/minute, compared to a reported speed of 1.5–6 µm/min in other cell types) when compared to passive organelle movement,^12,22,54^ potentially indicating active transport and likely involving motor proteins and actin filaments.^45,55^

We observed frequent contact between fibroblast NT and ECM. The exact relevance of this association is unclear. It is plausible that NT are involved in controlling ECM architecture, such as the placement of fibres or their cross-linking. In previous studies, integrins (which are key ECM-binding surface receptors) were reported to accumulate in filopodia, and have been suggested to prime NT to probe the matrix, creating ‘sticky fingers’ along the leading edge of the cell.^10,56^ The association of NT with ECM could also aid the cells in the process of ‘guidance’ or pathfinding through the tissue – this could be of potential relevance in fibrotic, ever-remodelling cardiac tissue during disease.^57,58^

Most of our studies were performed *in vitro*. Translating *in vitro* NT characteristics to possible *in vivo* relevance is challenging due to the morphological changes that occur in cultured fibroblasts, such as flattening and changes in the expression of fibroblast activation markers.^4,29^ In addition, the environment of the tissue would offer unique challenges to NT progression – NT would likely have to contort their shape due to obstacles, such as other cells and dense ECM, preventing the NT from connecting at the shortest distance between cells.^16,21^ While we observed NT-like structures in intact human atrial tissue, observing their dynamic behaviour in a 3D environment is technically challenging and would require extensive development of novel imaging methods.

Our study is potentially affected by a number of limitations. As the exact criteria for distinguishing different NT populations are not clear, we cannot differentiate between filopodia or tunnelling nanotubes for example, each of which could have vastly different roles, potentially confounding our results. We also performed most of our experiments in monoculture, and in future the inclusion of other cell types (such as cardiomyocytes or endothelial cells)^23^ would potentially provide further valuable insights into the relevance of NT-mediated heterocellular communication. In our intact tissue studies, we indeed observed regular arrays of electron-dense proteins at the point of contact between fibroblast NT and cardiomyocytes, suggesting that NT can interact with target cells via complementary surface cues, such as receptors and/or ligands. Additional experiments could explore the association with NT structure and their lifespan,^16^ the role of mechanosensors in the initiation of NT formation,^59^ additional types of the cargo that can be transported along NT (lysosomes, lipid droplets),^60^ and the possible mechanisms underlying the establishment of open-ended connections.^61^

Our results advance our understanding of how cardiac fibroblasts interact with their environment via NT. NT were shown to be very dynamic and labile structures, which supports their proposed role in rapid homo- and hetero-cellular communication, and their proposed role in enabling swift responses to changes in tissue state. NT properties and behaviour differed between fibroblasts derived from SR and AF tissue. We were able to alter NT structure and motility using pharmacological and environmental interventions. Furthermore, our observations indicate that NT play a role in the establishment and/or maintenance of the ECM matrix, highlighting them as very versatile structures capable of both sensory and actuating functions. The presence of NT-based communication in the heart would lend it ‘super-cellularity’ properties,^8^ allowing for rapid balancing of metabolic needs, stress responses, and ECM homeostasis through long-distance exchanges between cells that are not in immediate contact. In the heart, NT-based communication could also mediate the electrical coupling (also hetero-cellular)^3^ between distant cells; this is potentially accomplished with the aid of gap junction proteins,^62^ through the propagation of calcium transients, or through membrane depolarisation – a process that could be of crucial importance during pathological remodelling.^19,63,64^ Finally, more detailed knowledge of NT in the heart could offer novel ways of modifying specific cell properties, e.g. as a novel delivery route for drugs.^54^

## Acknowledgements

This work was supported by German Research Foundation Collaborative Research Centre SFB1425 (DFG #422681845) and Germany’s Excellence Strategy (CIBSS - EXC 2189). We thank the SCI-MED imaging facility at the Institute for Experimental Cardiovascular Medicine in Freiburg; Sarah Zimmermann, Ash Weier, and Tamara Basta at the CU Boulder Center for Cryo Electron Tomography; Martin Schorb, Rachel Mellwig, and the Electron Microscopy Core Facility at EMBL Heidelberg; Martin Helmstädter and the Electron Microscopy Core Facility at IMITATE in Freiburg; and Zoltan Metlagel (Thermo Scientific). ERZ is a German Research Foundation Emmy Noether Fellow (DFG #396913060). SSH, JG, AF, JM, JH, PI, MK, KK, CW, SN, TK, PB, RP, AR, PK, and ERZ are members of the German Research Foundation Collaborative Research Centre SFB1425 (DFG #422681845). Cryo-ET work was supported by the National Institutes of Health (NIH) Common Fund.

## Supplemental Figures

**Supplemental Figure S1.**
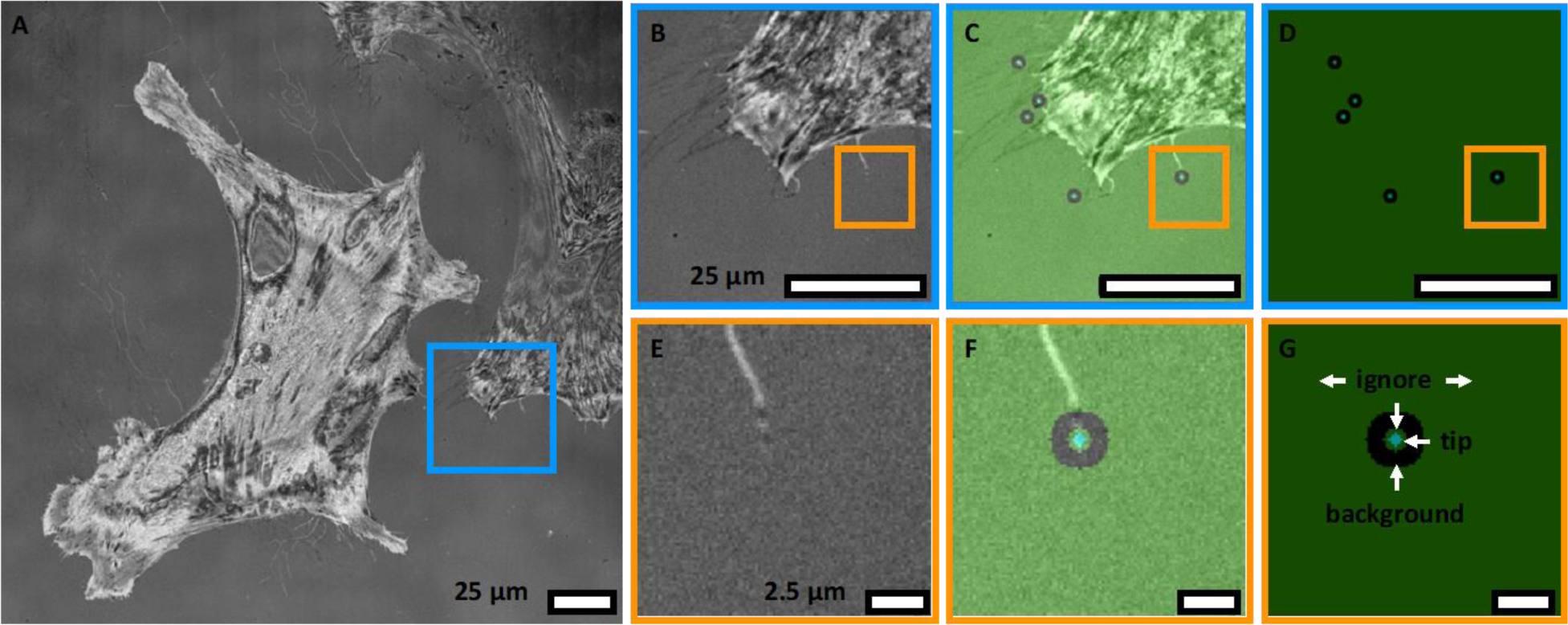
Sparse annotation strategy of NT. **A:** Typical reflection microscopy image showing fibroblasts with multiple NT. **B–D:** Magnified subsets of A, showing reflection image, overlay of reflection and annotation, and annotation, respectively. **E–G:** Magnified subsets of B, C, D, respectively. Sparse annotation strategy is shown in G: Sparse annotation strategy: the tip is annotated as foreground class (radius 2 pixels), surrounded by ignore class (2 pixels), surrounded by the background class (6 pixels).

**Supplemental Figure S2.**
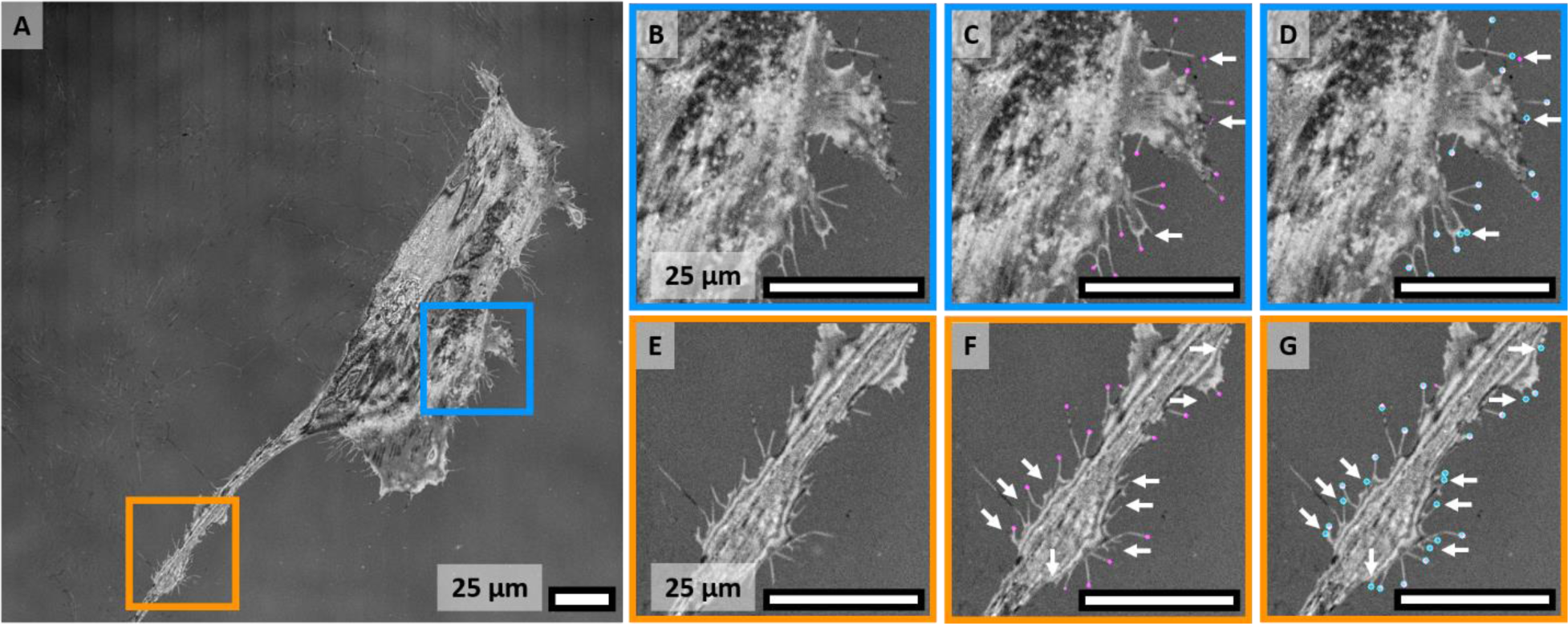
Deep Learning-based detection of NT tips. **A:** Reflection microscopy image from the test dataset. **B–D** and **E–G:** Magnified subsets of A, showing reflection image, prediction of the convolutional neural network, and overlay with annotation, respectively. White arrows indicate detection errors, predominantly appearing at short NT. B, C, D shows a subset with representative segmentation results; E, F, G shows a subset with particularly high error rates. The test dataset achieved a Dice coefficient score of 0.64.

**Supplemental Figure S3.**
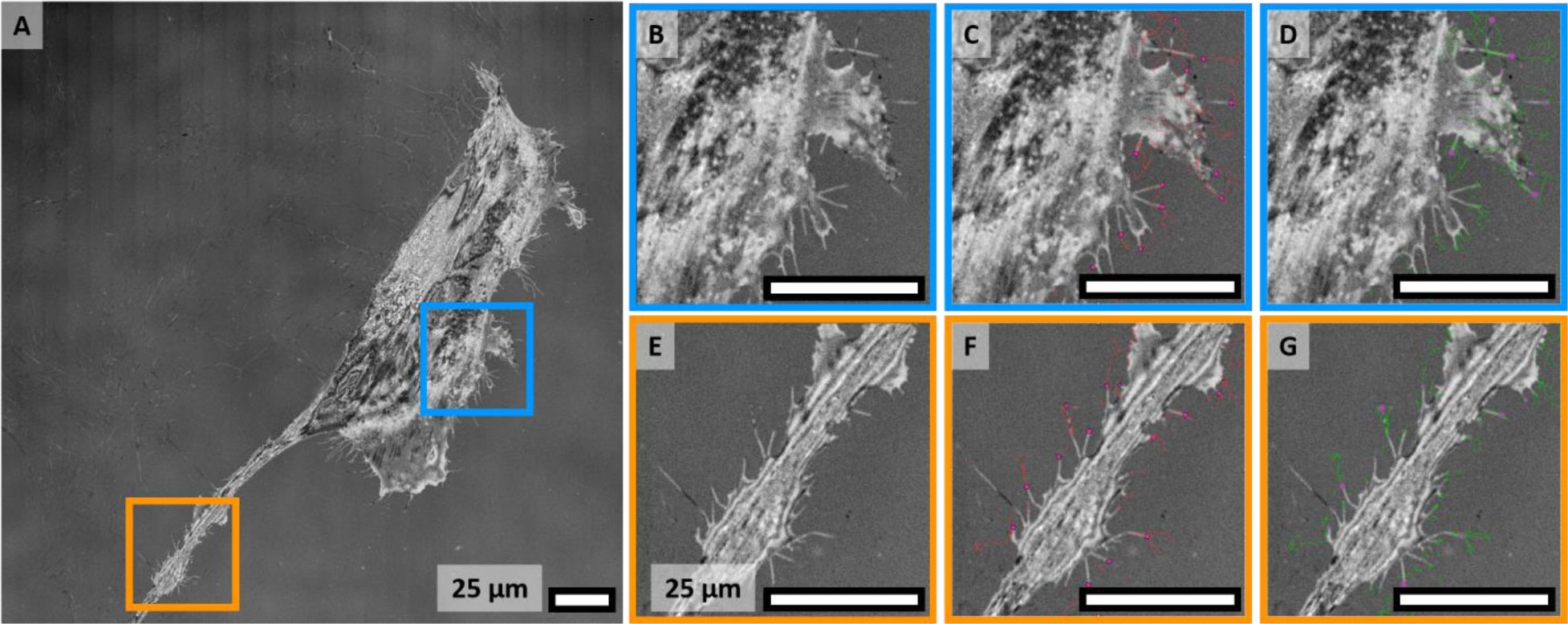
Tracking of automatically detected NT tips. **A:** Reflection microscopy image from the test dataset. **B–D** and **E–G:** Magnified subsets of A, showing reflection image, calculated tracks based on the predicted spots, and overlay with annotation, respectively. E, F, G shows a subset with particularly high error rates in the spot detection spot. The tracks on the test dataset resulted in a F1 score of 0.76.

**Supplemental Figure S4.**
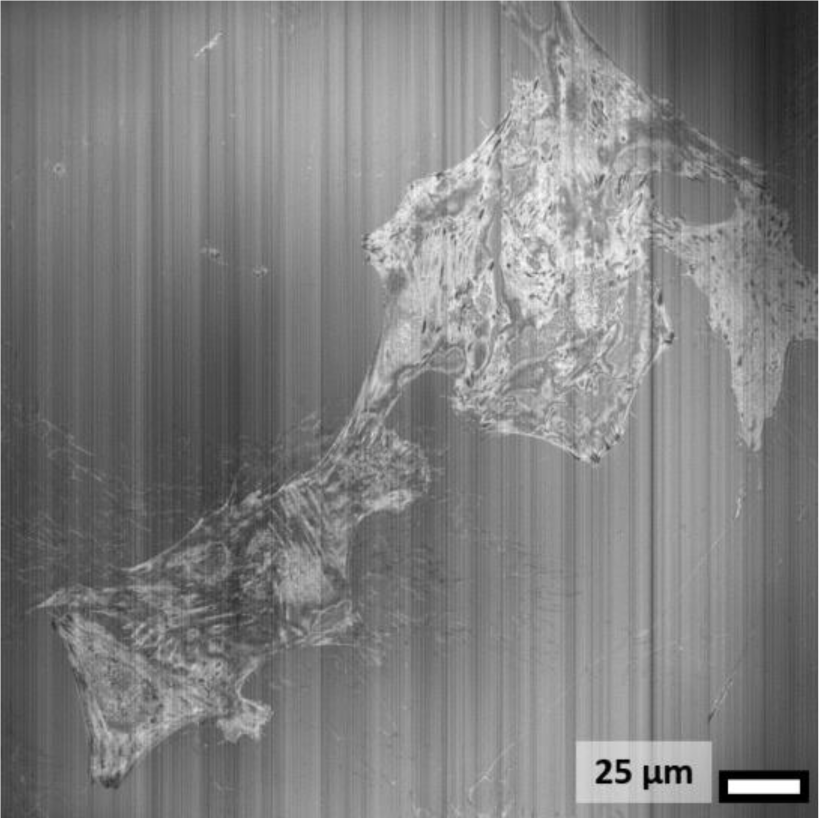
Representative example of excluded image stacks for NT motility analysis. The origin of this artefact is unknown.

**Supplemental Figure S5.**
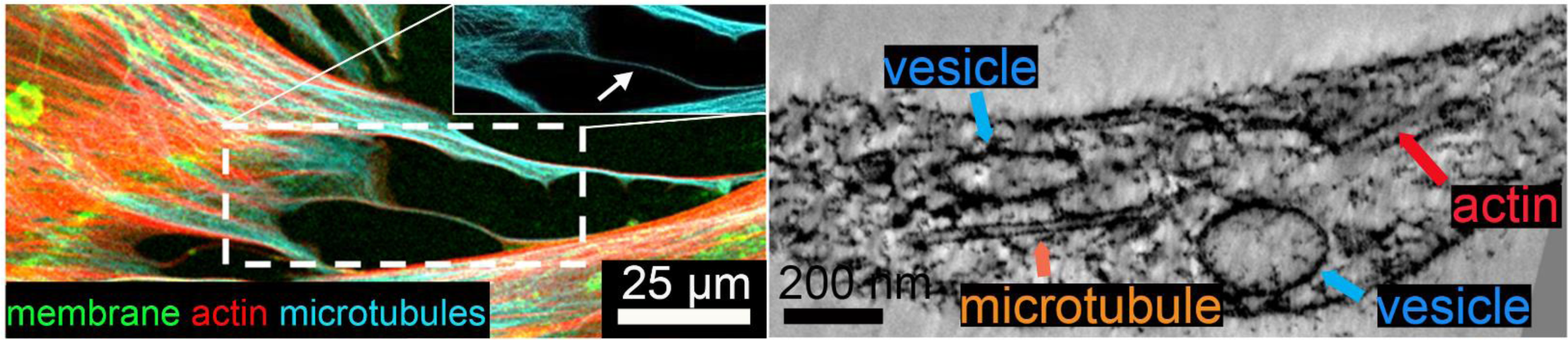
Microtubules in cardiac fibroblast NT. Microtubules could be occasionally identified using confocal microscopy (left, white arrow) or room temperature-ET of chemically-fixed cells.

## Abbreviations

AF: atrial fibrillation
ECM: extracellular matrix
ET: electron tomography
HAF: human atrial fibroblast (cell line)
NT: nanotubes
PBS: phosphate buffered saline
SR: sinus rhythm
TGF-β: transforming growth factor β

## Notes

### Competing Interest Statement

The authors have declared no competing interest.

### Summary of Updates

This version has been revised to correct the abbreviations in the abstract.

