## Supplemental Figures for "Structure and dynamics of human cardiac fibroblast nanotubes"

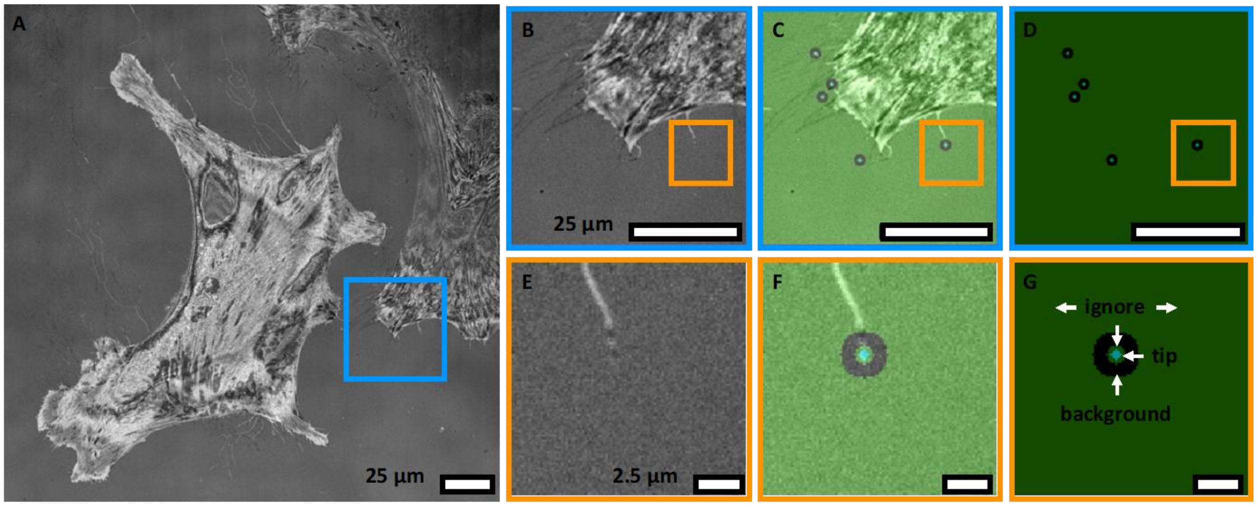


**Supplemental Figure S1. Sparse annotation strategy of NT. A:** Typical reflection microscopy image showing fibroblasts with multiple NT. **B–D:** Magnified subsets of A, showing reflection image, overlay of reflection and annotation, and annotation, respectively. **E–G:** Magnified subsets of B, C, D, respectively. Sparse annotation strategy is shown in G: Sparse annotation strategy: the tip is annotated as foreground class (radius 2 pixels), surrounded by ignore class (2 pixels), surrounded by the background class (6 pixels).

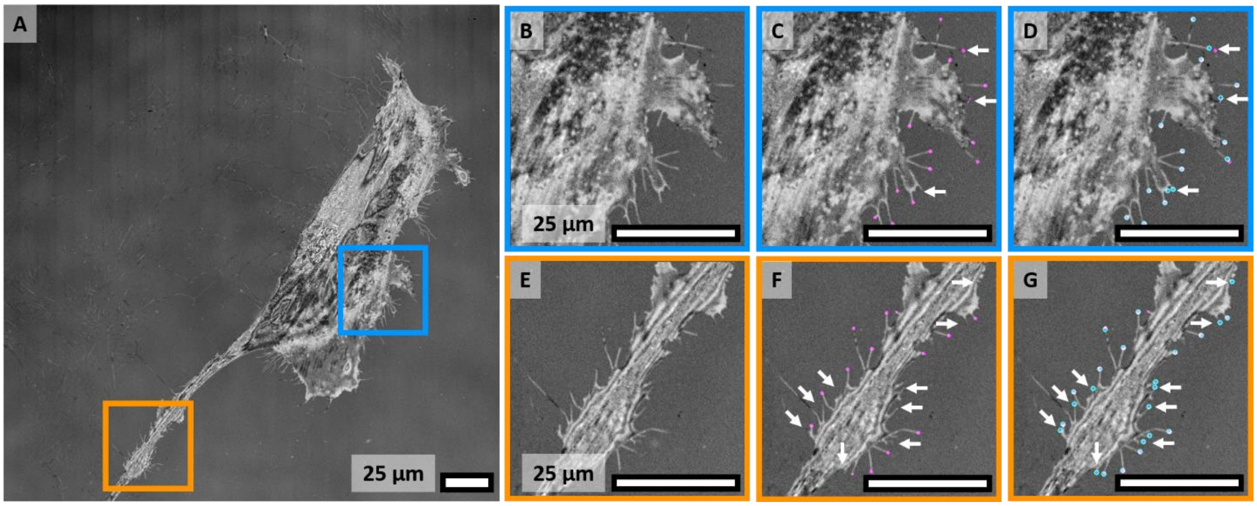


**Supplemental Figure S2. Deep Learning-based detection of NT tips. A:** Reflection microscopy image from the test dataset. **B–D** and **E–G:** Magnified subsets of A, showing reflection image, prediction of the convolutional neural network, and overlay with annotation, respectively. White arrows indicate detection errors, predominantly appearing at short NT. B, C, D shows a subset with representative segmentation results; E, F, G shows a subset with particularly high error rates. The test dataset achieved a Dice coefficient score of 0.64.

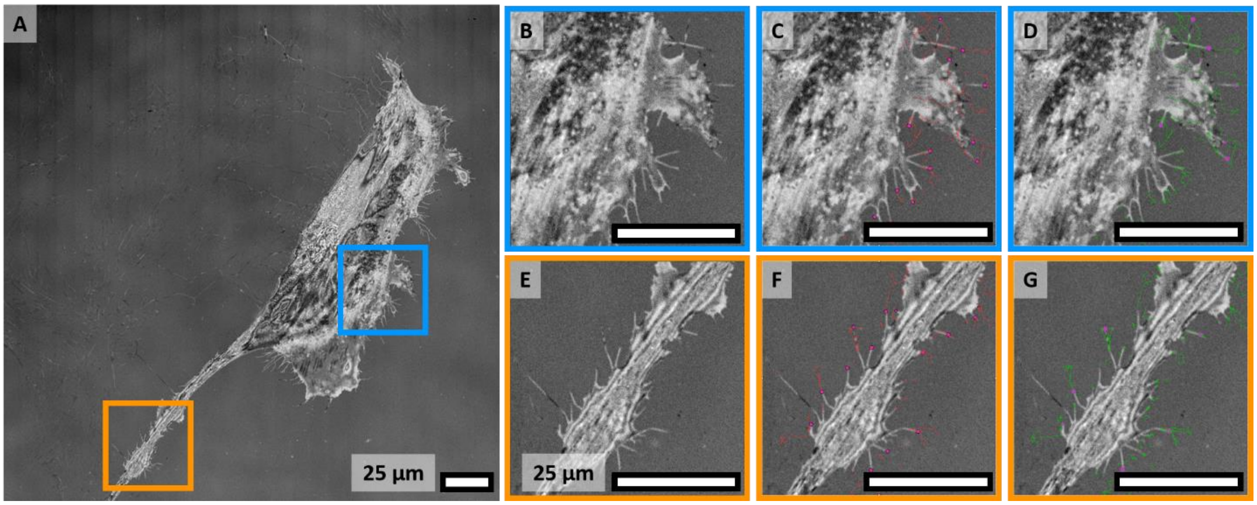


**Supplemental Figure S3. Tracking of automatically detected NT tips. A:** Reflection microscopy image from the test dataset. **B–D** and **E–G:** Magnified subsets of A, showing reflection image, calculated tracks based on the predicted spots, and overlay with annotation, respectively. E, F, G shows a subset with particularly high error rates in the spot detection spot. The tracks on the test dataset resulted in a F1 score of 0.76.

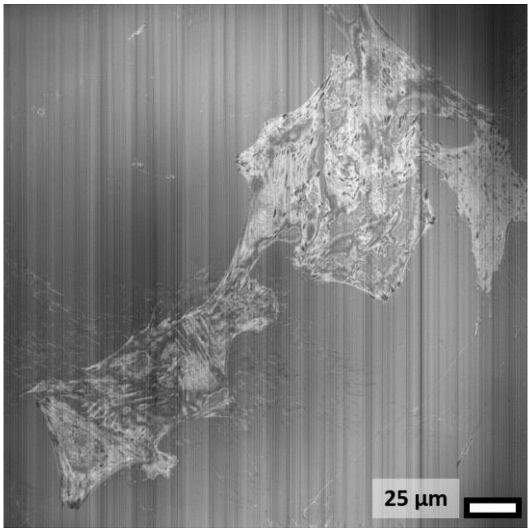


**Supplemental Figure S4. Representative example of excluded image stacks for NT motility analysis.** The origin of this artefact is unknown.

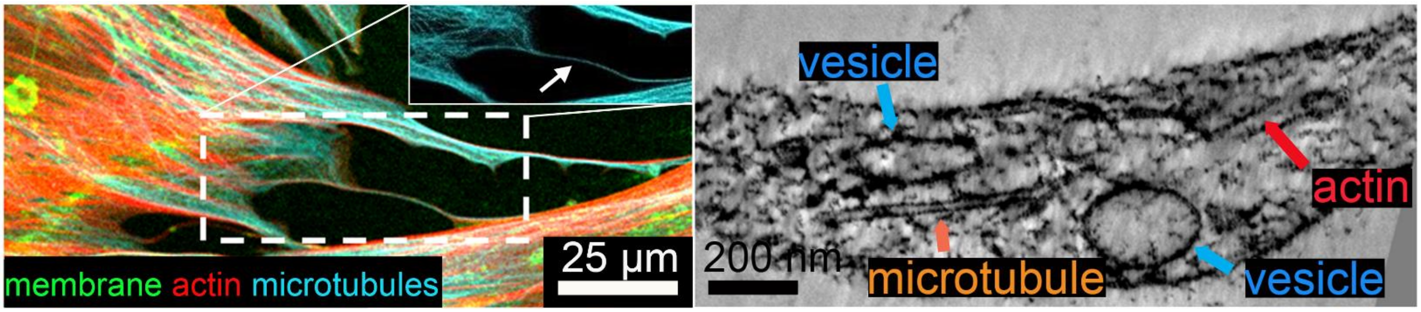


**Supplemental Figure S5. Microtubules in cardiac fibroblast NT.** Microtubules could be occasionally identified using confocal microscopy (left, white arrow) or room temperature-ET of chemically-fixed cells.
